# Wild sulphur-crested cockatoos match human activity rhythms to access food in the urban environment

**DOI:** 10.1101/2023.09.26.555651

**Authors:** G. Fehlmann, J. M. Martin, K. Safi, L. M. Aplin

## Abstract

Urban areas are growing rapidly across the globe, and wild species are occupying this new environment. Despite offering potential resources, disparities in the urban matrix can lead to specific challenges, with pathways and resources fragmented in space and time. Urban-dwelling species would therefore benefit from learning when and where to exploit human derived food. Here, we investigate whether birds synchronize the exploitation of the most urbanized areas to match food-provisioning patterns, using the example of the popular hand-feeding of sulphur-crested cockatoos (*Cacatua galerita*) in Sydney, Australia. We monitored the provisioning behaviour of people via a large-scale citizen science program, and tested for synchrony with the spatial behaviour of eight birds equipped with GPS loggers. Our data show that sulphur-crested cockatoos exploited the urban environment, relying on the green areas of the city; importantly, they also visited buildings within more urbanized areas. Sulphur-crested cockatoos used urban space with specific time patterns particularly matching human recreational feeding routines, suggestive of time-place learning. We show that urban environments provide daily temporal foraging resources for which species adjust behaviorally. Thus, our data support the general claim that retaining green spaces in cities is essential to sustainable urban planning, and key to allow species to exploit the urban environment, particularly in areas of high human density. This study builds on the literature investigating human-animal interactions, expanding our understanding of animals’ exploitation of human behavior. Further research should include the impact of such interactions on urban wildlife’s fitness according to their cognitive and behavioral traits.

## Introduction

Urbanization represents one of the most drastic changes that humans can impose on the environment (Vitousek et al., 1997). Yet a growing body of literature highlights how species have adapted to, and thrive in this highly modified environment (Fehlmann et al., 2021; Ritzel & Gallo, 2020; Sol et al., 2013). Indeed, the urban environment can offer new opportunities to wildlife, such as food, reduced climatic variation, and lower predation pressure (Shochat et al., 2006). As urban areas continue to grow rapidly around the globe, understanding how species exhibit behavioural adaptations that enable them to live successfully alongside humans is crucial for efforts to increase biodiversity in urban spaces.

For wildlife, urban adaptation is not a given (Fehlmann et al., 2021; Sol et al., 2013, 2014); finding space to sleep, forage and reproduce can be challenging due to the degradation of natural habitats, fragmentation, noise and light pollution, and disturbance by humans (Sol et al., 2014). City landscapes are divided in a mosaic of small properties individually managed by different landowners, leading to a lack of continuum in resources or pathways (Aronson et al., 2017). In addition to the physical environments, the presence of large populations of humans and the accumulation of individual routines result in large-scale flows of people, including high traffic when people commute to work and high frequentation of parks during weekends. Such patterns strongly shape the ecology of urban wildlife that adjust to these artificial rhythms. Shifts in diel activities are frequently reported in response to negative interactions with people (Ritzel & Gallo, 2020; Sol et al., 2013). For example, coyotes in Alberta, Canada, were found to be more likely to survive when active around midnight rather than at dusk leading to urban coyotes becoming more nocturnal (Murray & Clair, 2015). Yet, humans’ presence can also attract wildlife, in particular by creating foraging opportunities (for example household waste, outdoor lunches, deliberate provisioning) (Ducatez et al., 2013; Flint et al., 2016; Klump et al., 2021).

Human supplied foraging opportunities tend to be energetically rewarding, predictable in space and time, but can also be short-lived and dependent on human activity patterns (Barrett et al., 2018; Fehlmann et al., 2017; Klump et al., 2021). As a result, wildlife exploiting such resources can face time restrictions based on artificial cycles such as weeks or working schedules and it may therefore become advantageous to learn the time and place to benefit from human derived short-lived foraging opportunities (Barrett et al., 2018; Lee & Thornton, 2021). If so, such positive direct human-wildlife interactions can lead to synchronicity in activity, facilitating the co-existence of wildlife in urbanized areas. As many urban-dwelling species foraging on human provided resources are likely to be confronted with such tasks, knowing where and when species exploit the different environments within the urban matrix is essential.

Here, we use the example of human provisioning of sulphur-crested (SC) cockatoos (*Cacatua galerita*) in central Sydney, Australia to explore one such positive human-wildlife interaction (Kirksey et al., 2018). In contrast with most global bird feeding activity, local residents of Australian cities often feed birds ‘by hand’, either directly, or by placing handfuls of seed on balconies or windowsills in response to solicitation by visiting birds (Kirksey et al., 2018). SC cockatoos are a generalist and widespread parrot that are successful urban adaptors in cities across Australia, which in the city center of Sydney, have habituated to people to such an extent that they are regularly hand fed by local residents. SC cockatoos have been the subject of a long-running citizen science program since 2010, where people report sightings of 144 individually wing-tagged SC-cockatoos through a smart-phone application (‘Big City Birds App.’; Aplin et al., 2021; Davis et al., 2017). We equipped 8 wing-tagged SC cockatoos with GPS and matched this active tracking data with citizen-science data on human provisioning. We identified sites where birds would spend time and investigated which habitats the SC cockatoos exploited. We then explored the timing of the visits of the GPS tracked individuals to the different habitat types, hypothesizing that birds would synchronize their exploitation of the most urbanized areas to match the spatio-temporal patterning of human supplementary feeding.

## Methods

### Study population and area

We studied one population of sulphur-crested cockatoos (*Cacatua galerita*) in central Sydney. SC cockatoos are a large (700-1200g), slow breeding and long-lived parrot that forage on a variety of food sources including shoots, roots, seeds, nuts and fruits. In urban areas, their diet also includes bird seeds and nuts (e.g., almonds) voluntarily provisioned by people in parks or at their balconies. This is provided at traditional bird feeders, but often also provided by direct hand-feeding, making it a highly temporally restricted food source.

SC cockatoos form stable roosting groups of approximately 50-500 individuals that collectively exploit a home-range centered on this roost site (Penndorf et al., 2023). We focused on the two most central roosting groups, one in the Royal Botanic Garden (S -33.864377, E 151.214693, approx. 70 birds) and one in Clifton Gardens (S -33.837209, E 151.251112, approx. 100 birds), at 1.5 km and 5.3 km from Sydney’s center business district respectively. Both areas encompass trees and grass lands (mostly parks and private gardens) and built environments. Within a radius of 2.5km from the Botanic Garden, the land cover was characterized by 20.8% trees, 5.8% grass, and 62.6% built environment. Around Clifton Gardens, trees represent 44.1% of the land cover, 5.6% grass, and 43.73% the built environment.

We equipped 8 birds with solar powered GPS receivers (e-Obs GmbH), recording GPS locations every 5min in 2016 and 2017 from 5:00 to 21:00 local time. This resulted in three tagged birds roosting in Clifton gardens, and five from the Botanic Garden area. Data acquisition was uneven across the year and limited by solar recharge. We therefore focused our analysis on spring-summer (September-February), when the tags had the higher performance rate with an average of 61 (+/-45) fixes a day (min = 5, max = 169).

### Quantifying bird feeding activities

We documented recreational feeding of SC cockatoos using the citizen science project called ‘Wingtags’ (Davis et al., 2017). Launched in 2012, this project encourages local Sydney residents to report observations of wing-tagged SC cockatoos through a smart-device application, submitting a photograph of the observed bird(s) along with their identity (defined by a wing-tag number). The application then adds a location and timestamp to the observation.

The dataset available for this study included >23,000 reports across Sydney. Unfortunately, we could not retrieve photographs from March 2014 to January 2017 due to a data error. The remaining dataset included 13,515 reports that had attached photographs and were from within 3.5km of one of the two roosting sites. Of these photographs, 2,344 had been annotated by volunteers as part of the Australian Museum ‘DIGIVOL’ scheme (https://australianmuseum.net.au/digivol), recording the behavior of the bird in the picture. For the purposes of this study, we pre-screened all the remaining 11,170 photographs to identify potential foraging events. That is, if the bird was holding a foot up, was leaning forward, standing next to a feeder, on a person or inside a building. This resulted in 2,902 reports. We annotated photographs using the same scheme as the ‘DigiVol’ dataset used. We considered birds being fed by people if the bird was observed eating, holding or standing next to manufactured food items, nuts or loose seeds. Importantly, we did not include reports when birds were standing next to a bird feeder, focusing only on active recreational feeding. Since people can submit several reports for the same feeding event (when several birds were foraging), we only considered feeding events reported by the same person if they were more than 20 minutes apart. This resulted in 1657 reports between April 2012 and July 2021. We assigned each feeding event to either the Botanic Garden area or the Clifton Gardens area according to the distance to each roosting site.

### Space use

#### Definition of exploited areas

We used first-passage time (FPT) to identify areas that were exploited more intensely within the home range. Traditional FPT studies are based upon the assumption that foraging animals engage in more tortuous and slower movements (Area of Restricted Search: ARS, (Fauchald & Tveraa, 2003)), resulting in animals staying within a certain radius longer than when traveling. We used this method to identify where birds were stationary considering that the GPS signal of a stationary bird will result in a local Brownian movement pattern within the range of GPS error (around 30m). To meet FPT assumptions of regular sampling, we standardized GPS locations to a regular time lag of 5 minutes by linearly interpolating location in time for locations with less than 30 minutes time lag. When time gaps were larger than 30 minutes, we split the track for each individual into distinctive bursts and discarded all burst that lasted less than 1.5h.

We identified distinctive phases in the birds’ movements, i.e. sedentary vs travelling behaviours, by plotting FPT against time. We used the Lavielle segmentation to partition each burst (Barraquand & Benhamou, 2008). This segmentation process is used to identify the location and the number of change points in the time series data, breaking the signal in bouts of homogeneous means and variances (Lavielle, 2005). For each segment, we identified the location of the maximum FPT, where the animal stayed longest. The location was considered as a potential exploited site if FPT was above 30 min and below 6h.

#### Recursive visits

Most ARS resulting from the above analysis were clustered into small patches, suggesting recursive visits to the same site. In order to estimate the location of these recursively visited sites, we projected the location of each ARS identified on to a raster with a 30 m grid cell. We then iteratively considered local maxima to identify clusters of ARS. Within each iteration, we selected the cell containing the most ARS and its 8 neighboring cells. We averaged the location of all ARS contained within these nine cells and added a buffer of 45m around this point to account for GPS error. Finally, we considered all ARS falling within this circle as revisits to the same site, we removed these from the list of ARS before starting a new iteration. Sites that were visited less than 3 times over the entire duration of the spring and summer were not considered for later analysis.

#### Environmental sampling

We described the urban environment with a 2m resolution land cover map obtained in 2019 from the publicly available dataset “Geoscape” (PSMA). This dataset is based on satellite imagery, details on surface cover such as roads, buildings, built-up areas, swimming pool, bare earth, grass, trees, low vegetation, and water. We combined ‘roads’ and ‘built up areas’ as ‘urban features’, including roads, parking lots made of man-made substrate, built up areas smaller than 9m^2^, and other man-made environments. For each revisited site, we calculated the proportion of the area covered by grass, trees, low vegetation, buildings, urban features, and assigned to each site the dominant habitat feature. When buildings were present, we calculated the average building height and area and the proportion of residential buildings located at each site. In order to assign an urbanization index to each visited site, we performed a Principal Component Analysis including all the above-mentioned habitat data. We used the first component described by this analysis as an index of urbanization, negative values, indicating greener areas and positive values more urbanized areas (Fig. 3).

**Fig. 2.**
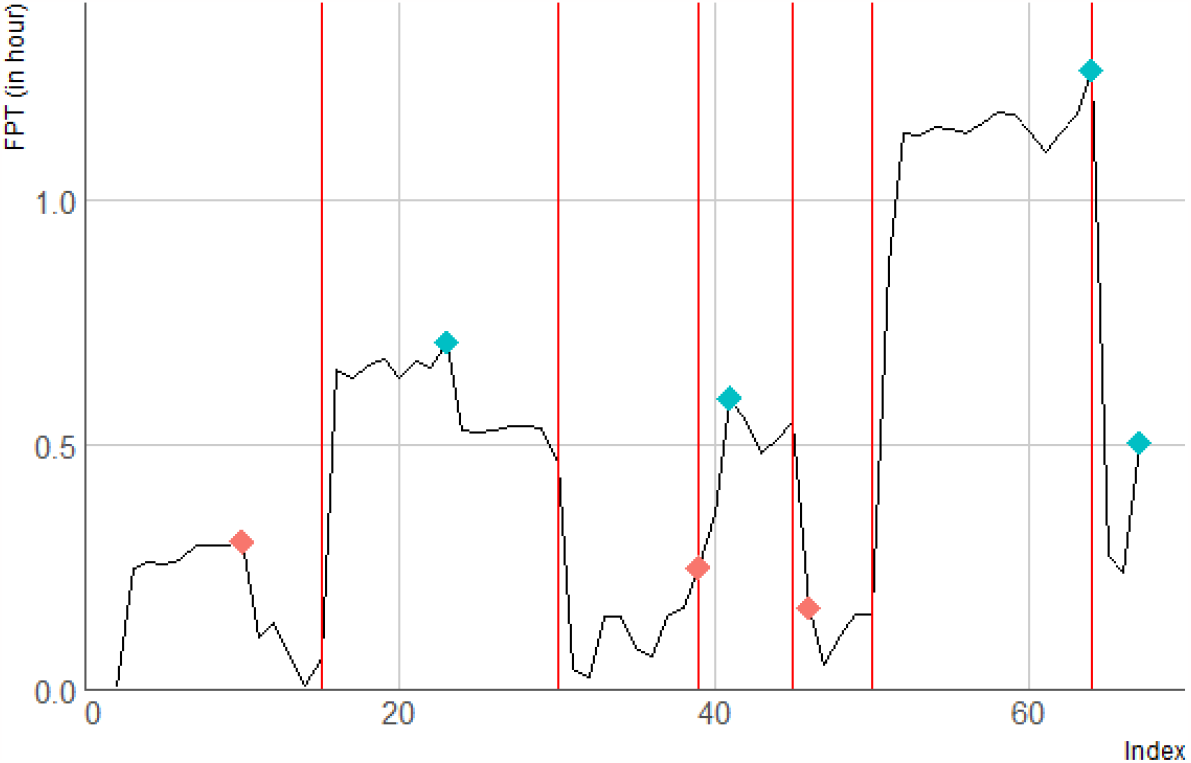
First Passage Time (in hours; FPT) plotted against time (as index) for one GPS burst. The result of Lavielle segmentation is represented with vertical red lines. Within each segment, we identified the locations at which the maximum FPT occurred, i.e. where the animal stayed the longest, and selected it as an Area of Restricted Search if FPT was higher than 30min and below 6 hours.

**Fig. 3:**
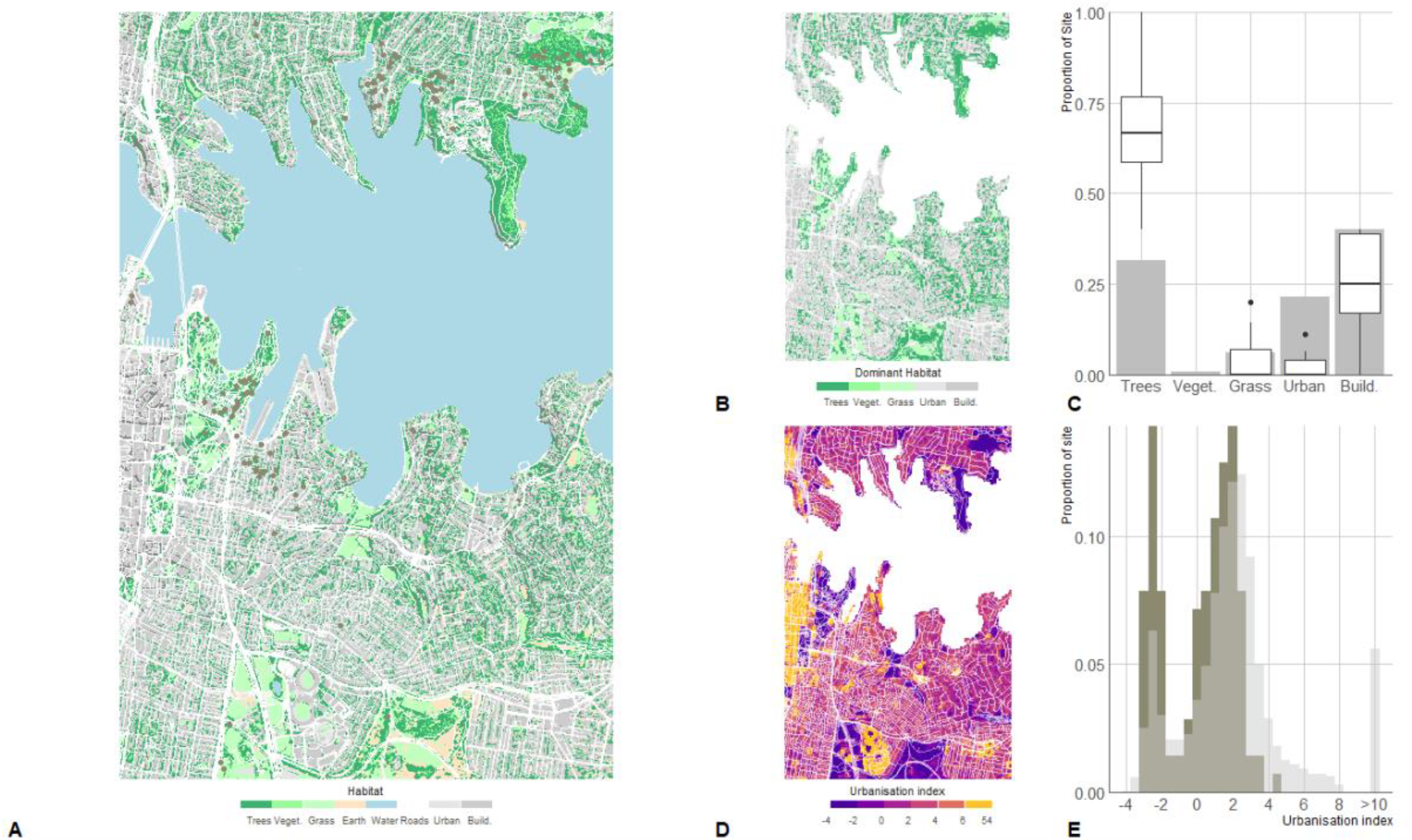
(A) The SC cockatoos equipped with a GPS (N=8) in the city center of Sydney revisited specific locations indicated by grey points. (B) We characterized these sites and the study area by considering the dominant habitat within 30x30m cells, and (C) compared each birds’ use of the different habitats (boxplot) versus the availability of habitats in the study area (bars). We investigated the effect of urbanization on birds’ space use by assigning an urbanization index score based on the proportion of the different habitat features, building size, height and use using Principal Component Analysis (D). We then explored the distribution of the exploited sites (except the roosting sites) across the urbanization index (dark grey bars) in comparison to the classification of the study area (light grey bars) (E). In B and D, roads (in white) are only given as visual landmarks.

#### Statistical analysis

We compared the timings of bird visits to buildings versus the timing of their visits to any of the other habitat types using a logistic regression. For each visit to an exploited site (ARS), as indicated by the GPS tagged birds, we used the habitat classified as 1 for buildings and 0 for any of the other habitat type as our response variable. We included the date, the hour of the day, the weekday and the month as predictive variables and tested for any interactions between these terms and the roosting site of each bird (Botanic Garden or Clifton Garden). We controlled for individual effects by adding individual as a fixed effect term in the model. We tested the value of each term independently and selected the best model according to AIC. In order to avoid sampling bias we performed 100 iterations, sampling randomly 300 visits at buildings and 300 visits to other habitat types and selected the most common winning model (based on AIC) as our final selected model.

We then described the timing of visits to buildings. We created time series with one-hour resolution from 06:00 to 20:00, only including timestamps for which GPS locations were recorded. For each time stamp, we reported the number of visits per bird. Time was reported in AEDT (UTC + 11) throughout the study period. We used General Additive Models (GAM) in R (R Core Team, 2019) using the mgcv package (Wood et al., 2016) with a zero inflated Poisson distribution to fit the seasonality of visits considering daily, weekly and monthly patterns. For modelling daily and monthly variations, we used cubic spline functions allowing a maximum of 16 and 6 knots respectively (because GPS were recording data for 15 hour/day and 6 month/year). For weekly variations, we used cyclic cubic splines setting a maximum of 7 knots. We allowed the model to compute a penalty for each smooth term to simplify their ‘wiggliness’ and reduce their effective degrees of freedom (Wood et al., 2016). We tested the significance of interactions between hours and weekdays and hours and months by adding tensor product smooths. To account for site variations between the two roosts, we allowed different seasonal patterns at each site. Because our recording encompassed daylight saving time change, we considered its potential impact in hourly variations by computing separate smooth terms, before and after time change. We controlled for individual variation and year effect by adding these as fixed effects and for accounting for temporal autocorrelation and sampling variations by adding the timestamp as smooth term using p-splines and a maximum of 24 knots.

To document human provisioning, we identified in which habitat it occurred via classification of the pictures and assigned an urbanization index value to each location using our PCA model. We then tested with a Wilcoxon rank sum test the differences in urbanization index score for sites around the Botanic Gardens vs Clifton gardens. We discretized the temporal patterns in recreational feeding of birds by creating a time series of the number of bird feeding event to resolution of one-hour. For consistency between GPS and citizen science data, time was reported in AEDT (UTC + 11) throughout the year. We used GAM with a zero inflated Poisson distribution to model the seasonality of the behaviour taking into account monthly, weekly and daily patterns using cyclic cubic splines with a maximum of 12, 7 and 12 knots respectively. We tested for differences in human provisioning patterns in the two studied areas by allowing these monthly, weekly and daily patterns to vary according to site. We took into account daylight saving time by estimating separate smoothed terms for hourly patterns for each period (Standard time and Daylight Saving Time). We also tested for the interaction between daily and weekly patterns and between daily and monthly patterns using a tensor product smooth. We controlled for temporal autocorrelation by adding the timestamp as smooth term using p-splines and a maximum of 24 knots (for the 24 months of data).

To test for synchrony between birds’ foraging patterns and human provisioning, we predicted bird visits to buildings and recreational feeding from October 2017 to March 2018 around Clifton Gardens and the Botanic Garden using our GAM models. We then used a partial mantel test, using the package ‘vegan’ (Oksanen et al., 2014) with 1 000 permutations to compare the variations of these two variables through time (hour and month).

## Results

### Recursively exploited sites

We identified 4495 ARS, indicating stationary behaviors for all birds. Of these ARS, 93.4% (± 6.3) were located within 45m of at least three other ARS, indicating 188 re-visited sites. Among these re-visited sites, the roosting site itself contained on average 53.2% (± 27.3) of the total ARS identified for each bird. Roosting sites were located in parks with remnant old-growth eucalyptus trees (min - max tree coverage: 46% - 75%) with a low urbanization index (min: -3.1, max: 0.9). As a reference, cells with such environmental characteristics (minimum of 46% tree coverage and urbanization index value below 0.9) represented 19.53% of the grid cell within the area.

Birds recursively exploited 181 other sites. These were on average within a distance of 921m from the birds’ roosts (sd: +/-780m). Among these sites, 68.0% (± 17.0) were covered mostly by trees, 25.3% (± 13.4) mostly by buildings, 4.4% (± 7.0) mostly by grass, and 2.3% (± 3.7) mostly by other urban features. As a comparison, the study area is covered by 31.5% of trees, 40.1% buildings, 6.0% grass, 21.5% urban features, and 0.9% low vegetation (excluding the surface covered by the sea, Fig. 3). We identified 486 visits to buildings, 84.4% of which occurred in residential areas (sites covered by more than 50% of residential buildings). All visits to non-residential buildings occurred within the Botanic Garden area where visits to non-residential areas represented 30.6% of all visits to buildings. Most recursively visited sites occurred in areas with low urbanization index scores (score below 0: 47.9%) or areas with median scores (score between 0 and 3: 48.4%). Only 3.7% of revisited sites occurred in areas with an urbanization index above 3, despite this category representing 24.9% of the cells in the study area (Fig. 3).

### Timing of visits

Within our sampling schedule (from 5:00 to 21:00), roosting sites and trees were predominantly used during the morning and around midday, with no specific weekly patterns. Buildings were visited by the tracked birds from 8:00 to 18:00 (GAM: edf = 8.35, Ref.df = 14, chi^2^ = 54.87, p < 0.001), and significantly later than visits to other habitat types (Fig. 4 a. Logistic regression: estimate = 0.13, s.e. = 0.03, z = 4.6, p < 0.001). Visits to buildings were more frequent during weekdays, particularly Tuesdays and Thursdays (Fig. 3 b. Logistic regression: estimate = -0.17, s.e. = 0.05, z = -3.4, p < 0.001). Weekly fluctuations were however not significant when modelling temporal patterns across the two roosting areas (GAM: edf = 0.00, Ref.df = 5, chi^2^ = 0.00, p = 0.698).

**Fig. 4:**
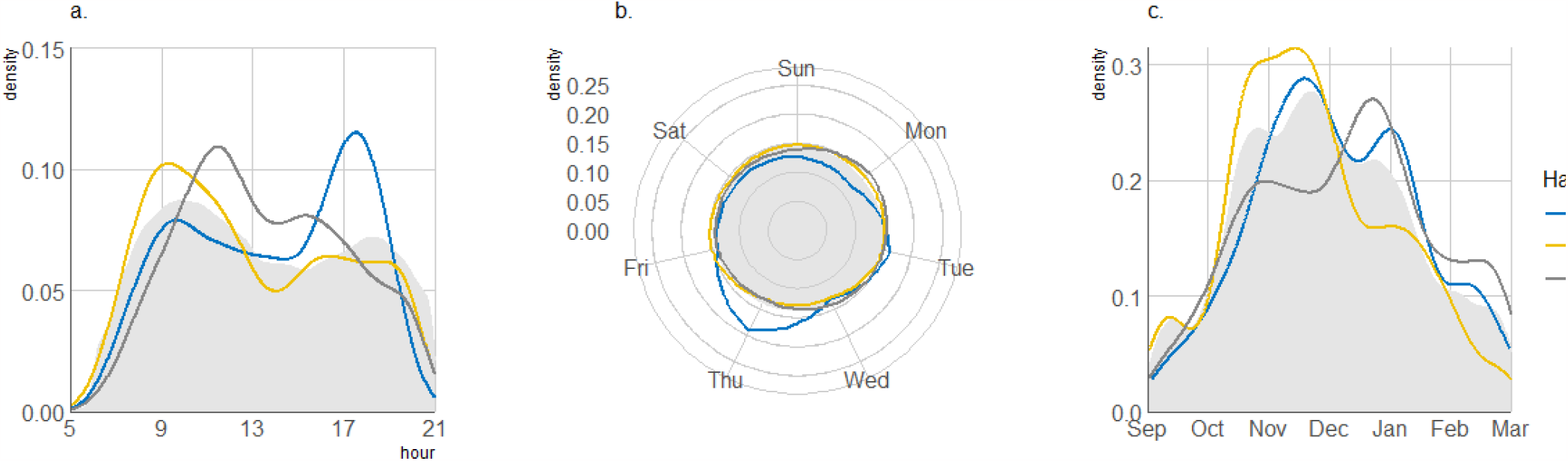
Time differences between visits to roost, buildings and trees considering (a.) hour of the day, (b.) day of the week, and (c.) month. Note that GPS recording depended upon solar input. Locations were recorded from 5:00 to 21:00 and data from March to September were not included in the study due to too few recordings. The density of GPS location through time is represented by the light grey area.

Across the study area, roosting sites were used most intensively during November, which coincided with nesting and chick rearing. Buildings were visited with a specific monthly pattern varying according to the birds’ ranging area (Fig.4 c. Logistic regression: BG: estimate = 1.3, s.e = 0.23, z = 5.70, p < 0.001, CG: estimate = 0.25, s.e = 0.07, z = 3.33, p < 0.001). Buildings located around the Botanic Garden were visited around 18:00 from September to December and throughout daytime in January. Buildings located around the Clifton Gardens were more specifically visited around 18:00 during October-December and February and less frequently visited in January (Fig. 4; GAM tensor (hour, month, by site): BG: edf = 11.70, reference df = 89, chi^2^ = 54.87, p < 0.001, CG: edf = 10.69, reference df = 89, chi^2^ = 31.12, p < 0.001).

The timing of visits to buildings did not vary through the day despite daylight saving schemes (GAM without vs with daylight savings: difference in degrees of freedom = -10.0, difference in AIC = 10.88). Buildings’ visitation rate also depended on time autocorrelation and sampling variation (edf = 1.17, reference df = 23, chi^2^ = 7.99, p = 0.003), and bird’s individual patterns (GAM: df = f, chi^2^ = 23.99, p < 0.001).

### Recreational bird feeding

Across the study area, 120 people participating in the Big City Birds Project actively fed and reported individually wing-tagged SC cockatoos (56 around the Botanic Garden and 65 around Clifton Gardens) throughout all months of the year. Most people fed birds from residential areas with 74% of reports (versus to 1.9% in grass and 0.3% in trees), which could be separated, to balconies (51.8%), windowsills (20.0%), or inside buildings (3.2%). Reports were mostly located in areas with an urbanization score comprised between: 1.6 – 2.9 (1^st^ – 3^rd^ quartile). Participants around Clifton Gardens reported feeding birds in significantly greener areas than around the Botanic Garden (CG median = 1.7, BG median = 2.7, Wilcoxon test: W= 547865, p<0.001).

Feeding occurred with a monthly and daily pattern, which varied according to the location (GAM; tensor (hour, month, by site): BG edf = 38.58, reference df = 120, chi^2^ = 146.4, p < 0.001; CG edf = 27.20, reference df = 120, chi2 = 191.7, p < 0.001, see Fig. 5). Most reports occurred at 18:00 throughout the year with a peak from September to March. A second peak of feeding occurred in the morning at 8:00 around the Botanic Garden but not Clifton Gardens. Most reports occurred from 9:00 to 18:00 (AEDT) during Australian Eastern Standard Time (UTC + 10) and from 8:00 to 18:00 (AEDT) during Australian Eastern Daylight Time (UTC + 11, GAM hour by time zone: AEST: edf = 8.57, reference df = 10.0, chi^2^ = 123.3, p < 0.001; AEDT: edf = 8.81, reference df = 10.0, chi^2^ = 151. 5, p < 0.001). Note that, for consistency, we modelled time in AEDT all year round, considering civil clock people therefore reported feeding birds from 8:00 to 17:00 during AEST and from 8:00 to 18:00 during AEDT. The number of reports collected also varied daily (GAM: edf = 8.58, reference df = 8.92, chi2 = 293.0, p < 0.001).

**Fig. 4a:**
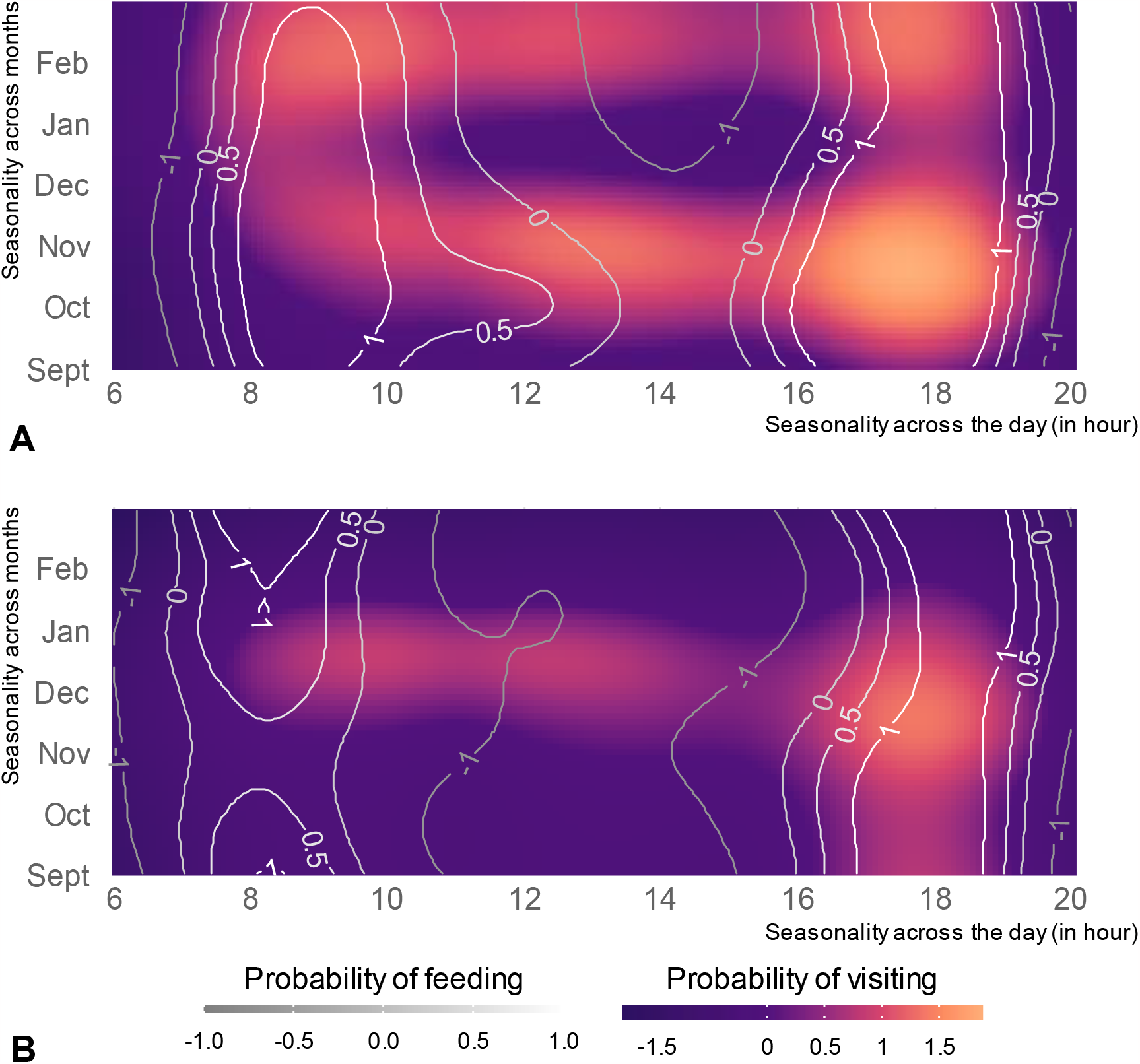
Birds fitted with GPS at Clifton Gardens (A) and Botanic Garden (B) visited buildings with distinctive time patterns over the day and throughout the month (color scheme). We compare this pattern to the recreational feeding patterns (contour plot), that was also following variations throughout the day and the month. The likelihood/ of bird visits was predicted by the following General Additive Model: probability of visiting ∼ (Hour:Month, by Site) + Hour + Date + Individual) and people fed bird according to the following model: probability of feeding ∼ (Hour:Month, by Site) + (Hour, daylight saving) + Date. We focused our analysis from September to February (included), time during which GPS sampling was high (we control for data sampling in both model by including date as a smoothed factor). Note that we scaled predictions for the two models to allow comparisons.

The timing of visits of the GPS tagged birds to buildings was correlated with the timing of human provisioning at both sites, although the effect was stronger at Clifton Gardens. Clifton Gardens: Mantel statistic r = 0.49, p<0.001; Botanic Garden: Mantel statistic r: 0.11, p<0.001 (significance test based on 1000 permutations).

## Discussion

Our study shows that large parrots living in the city center of Sydney megalopolis can tolerate an urbanized landscape, exploiting greener areas with a high proportion of trees, and visiting urban features with specific time patterns correlated with human recreational feeding routines. This is in line with previous findings showing that SC cockatoo populations have increased over the past decades, and exhibit similar or higher abundance in suburban and urban areas than in the surrounding natural habitat (Burgin & Saunders, 2007; Davis et al., 2012). Yet, our study also suggests that there are limits to this tolerance, with our GPS birds avoiding areas where the urbanization index was high (above 3). This suggests that highly urbanized areas with tall and large buildings, mostly covered by infrastructure and only little vegetation, may not be suitable for SC cockatoos and other wildlife.

Our data support the general claim that retaining green spaces in cities is essential to sustainable urban planning (Campos-Silva & Piratelli, 2021), and key to allow species to exploit the urban environment. SC cockatoos did successfully exploit areas of medium urban density when foraging, however roosting sites and the most intensively exploited areas were greener, covered by at least 43% of trees and located in less urbanized areas such as parks or patches of remnant vegetation. When wildlife can tolerate urban features, or benefit from them, this frequently relies on adjacent patches of more natural vegetation as a refuge or for reproduction (Davis et al., 2013; Gallo & Fidino, 2018; Grafius et al., 2017). Land cover is one of the main factors influencing bird diversity in urban landscapes, with patches of remnant vegetation hosting more biodiverse birds communities (Aronson et al., 2014; Callaghan et al., 2018; Kuras et al., 2020).

Provisioning of wildlife, for example via bird feeders, is a globally popular activity (Reynolds et al., 2017). For example, in the high density Sydney central district, 2 to 3 people reported feeding SC cockatoos per km^2^, with most feeding occurring directly at people’s balconies or windowsills. Such sites were exploited at even higher rates than grass patches, foraging sites that form an important part of the natural repertoire of this species (Polley & Lill, 2021). In urban areas, parks are predominantly lawn monocultures, with low plant and animal biodiversity (Aronson et al., 2017), and ground foraging may be vulnerable to attack from domestic dogs and harassment from children. Balconies and windowsills may thus appear as a more attractive foraging space.

Furthermore, the nutritional reward of seeds and nuts may eclipse that of grass leaves, shoots, seeds, and roots. Research into the nutritional benefits of urban foraging resources is warranted, particularly in the context of cognitive and behavioral traits. For wildlife that can solicit or take advantage of provisioning, such behavioral adaptations can have large consequences for the structuring of urban bird assemblages (Callaghan et al., 2019; Fuller et al., 2008; Galbraith et al., 2017).

Our analysis of citizen science reports of such feeding of SC cockatoos showed that provisioning was cyclic, with most feeding reported either at 8:00 or between 17:00-18:00. While our data collection could not ascertain the causation of this patterning, the close match to the most common work hours strongly suggests that this was driven by human activities. The visits of the GPS tagged birds to buildings mirrored the peak of this feeding activity. Indeed, the timing of the 486 identified visits was restricted in time, with a more distinct pattern than visits to trees or the roost; they occurred later in the day, and were overall correlated with human provisioning. Other recent work also suggest that SC cockatoo are able to take advantage of periodic human activities, such as household bins, only available to SC cockatoos once a week when put on the curb for collection (Klump et al., 2021, 2022).

Episodic memory (what-when-where) has been shown in food-caching species, such as corvids, that are able to integrate spatial-temporal information to return to a cache before food items perish (Grodzinski & Clayton, 2010). Here, the process may be comparable and allow these birds to adaptively exploit specific sites in more urbanized areas. Interestingly, birds living in the most urbanized area (around Botanic Garden) and visiting residential and commercial/business buildings expressed a different daily rhythm which did not correlate as strongly to feeding patterns as birds at the inner suburb roost of Clifton Gardens that visited only residential buildings. Indeed, people feeding SC cockatoos from their office window would likely happen during business hours and therefore encourage birds to visit buildings during daytime. Unfortunately, visits to such areas were rare and did not allow us to model the periodic patterns of such visits. In addition, the Botanic Garden is also one of the most visited areas of Sydney (Hale & Macdonald, 2005) and even if feeding SC cockatoos is discouraged in the park, it is still often undertaken by visitors. Combined, these may break the clear diel rhythm of bird feeding observed in more residential areas. This could suggest more individual-specific foraging strategies, with more continuous sampling of multiple sites throughout the day; however, our sample size did not allow us to explore individual variation in more detail.

Our results highlight the unique opportunity that studies on urban wildlife and the ecosystems they are adapting to and exploiting have for understanding urban biodiversity establishment and maintenance, but also cognitive ecology. Such environments present specific challenges to wildlife, relatively recent in regard of species evolution time (Lee & Thornton, 2021). Here, we have combined direct tracking of an urban-adapted parrot to identify key resources, and combined this map with a citizen science approach to investigate human-wildlife interactions in the urban landscape. Our data suggest that SC cockatoos do not use all parts of their home range equally, but use green spaces as roosting and foraging areas, while facultatively using more urbanized areas at specific times when they were the most rewarding. This further implies a role for sophisticated time and place learning, with birds matching activity to human patterns. Parrots are amongst the most successful urban adaptors, with some urban populations even potentially acting as refuges for species that are declining and endangered in their natural habitat or range (Davis et al., 2012, 2013; de Matos Fragata et al., 2022). With such knowledge, we can improve planning to help urban areas become as suitable as possible for globally threatened bird taxa (Aronson et al., 2014; Major & Parsons, 2010; Old et al., 2014) and urban tolerant wildlife in general.

## Acknowledgements

We thank Adrian Davis and Richard Major for their role in establishing the wing-tag project and support to LMA. We are grateful to Elham Nourani, Anne Scharf, Julia Penndorf, Barbara Klump, Michael Chimiento for discussions, and Martin Wikelski for financial support. GF was co-funded by the Department of Biology of the University of Konstanz and the Ministerium für Wissenschaft, Forschung und Kunst via the Brigitte Schlieben Lange Programm. LMA was funded by a Max Planck Group Leader Fellowship, and

## Authors contribution

GF, LMA, and KS conceived the study. LMA and JM equipped the birds and collected data. GF processed the bird and citizen science data, and performed the analysis. GF wrote the manuscript with the support of KS. All authors commented previous versions of the manuscript, and read and approved the final manuscript.

*The authors have no relevant financial or non-financial interests to disclose*

